# Myosin waves and a mechanical asymmetry guide the oscillatory migration of *Drosophila* cardiac progenitors

**DOI:** 10.1101/2022.03.25.485827

**Authors:** Negar Balaghi, Gonca Erdemci-Tandogan, Christopher McFaul, Rodrigo Fernandez-Gonzalez

## Abstract

Heart development begins with the formation of a tube, as cardiac progenitors migrate from opposite sides of the embryo and meet medially. Defective cardiac progenitor movements cause congenital heart defects. However, the mechanisms of cell migration during early heart development remain poorly understood. We investigated the mechanisms of movement of the *Drosophila* cardiac progenitors, the cardioblasts. Using quantitative time-lapse microscopy, we found that cardioblasts did not advance monotonically. Instead, cardioblasts took periodic forward and backward steps as they migrated. The forward steps were greater in both amplitude and duration, resulting in net forward movement of the cells. The molecular motor non-muscle myosin II displayed an alternating pattern of localization between the leading and trailing ends of migrating cardioblasts, forming oscillatory waves that traversed the cells. The alternating pattern of myosin polarity was associated with the alternative contraction and relaxation of the leading and trailing edges of the cell. Mathematical modelling predicted that forward migration requires the presence of a stiff boundary at the trailing edge of the cardioblasts. Consistent with this, we found a supracellular actin cable at the trailing edge of the cardioblasts. When we reduced the tension sustained by the actin cable, the amplitude of the backward steps of cardioblasts increased, thus reducing the net forward speed of migration. Together, our results indicate that periodic cell shape changes coupled with a polarized actin cable produce asymmetrical forces that guide cardioblast migration.

## Introduction

Heart development in vertebrates and invertebrates begins with the formation of a primitive tube as cardiac progenitors migrate collectively from opposite sides of the embryo and meet medially (DeRuiter et al., 1992; Bodmer, 1995; Moorman and Christoffels, 2003). Improper formation of the early heart tube can lead to devastating congenital heart disorders, which remain the most common birth defects (Hoffman and Kaplan, 2002; Marelli et al., 2014; Irvine et al., 2015). While the gene regulatory networks that regulate cardiac cell fate specification are well characterized (Olson, 2006), little is known about the molecular mechanisms that govern the migration of cardiac progenitors, or the underlying cellular causes of many congenital heart defects.

In *Drosophila*, the embryonic heart is a linear structure, composed of 52 pairs of cardiac progenitors, the cardioblasts (Rugendorff et al., 1994; Bodmer, 1995). Cardioblasts form two organized columns, one on either side of the embryo, that migrate dorsally and medially to join their contralateral partners and form a tube. As cardioblasts migrate, they are accompanied by a group of cells at their trailing edge, the pericardial cells, that eventually contribute to the excretory system of the fly (Mills and King, 1965). Cardioblast movement occurs concurrently with dorsal closure, a process in which the epidermis spreads dorsally over an extraembryonic tissue, the amnioserosa (Kiehart et al., 2000). Throughout embryonic development, cardioblasts establish transient adherens junctions with the overlaying epidermis (Rugendorff et al., 1994; Tepass and Hartenstein, 1994). The strength of the adhesion between the heart and epidermis is greatly reduced as the cardioblasts approach the dorsal midline, when cardioblasts display autonomous migration independent of the overlaying tissue (Haack et al., 2014). The extent to which the overlaying epidermis contributes to cardioblast movement remains unclear.

In eukaryotic cells, networks formed by the cytoskeletal protein actin and the molecular motor non-muscle myosin II (hereafter referred to as myosin) produce forces critical for cell movement (Trepat et al., 2009). In *Drosophila* embryos, cardioblasts extend actin-rich protrusions that mediate contralateral partner recognition (Haack et al., 2014; Raza and Jacobs, 2016; Zhang et al., 2018). Myosin transiently accumulates at the leading edge of migrating cardioblasts, where it regulates protrusive activity (Vogler et al., 2014; Zhang et al., 2020). However, the role of actomyosin networks in cardioblast migration is not well understood.

## Results and discussion

### Cardioblasts oscillate back and forth during migration

To establish the mechanisms of cardioblast migration, we imaged embryos expressing mid^E19^:GFP (Jin et al., 2013), a cardioblast-specific nuclear marker, at stage 14 of embryonic development (approximately 11 hours old) (Fig. 1a, Movie S1). We used a support vector machine—a supervised machine learning classifier (Wang and Fernandez-Gonzalez, 2017; Fernandez-Gonzalez et al., 2021)—to detect cardioblast nuclei, and watershed-based segmentation coupled with particle image velocimetry (Wang et al., 2017) to delineate and track individual nuclei and reconstruct their trajectories (Figs. 1a’-a’’ and S1a-b, Movie S2). We found that the medial-lateral cell velocities were not constantly directed medially towards the dorsal midline, but rather oscillated between positive (medial) and negative (lateral) values (Fig. 1b-c). These results suggest that cardioblasts take forward (medial) and backward (lateral) steps as they migrate. On average, the period of oscillation was 2.34±0.03 minutes (mean±standard error of the mean, s.e.m.). The amplitude of the medially-directed steps was 2.13±0.02 times greater than the amplitude of laterally directed steps *(P* < 0.0001, Fig. 1d), and medially-directed steps lasted 1.76±0.02 times longer (*P* < 0.0001, Fig. 1e), thus contributing to the net medial movement of the cells. We obtained similar results when we quantified cardioblast movement in embryos expressing Dlg1:GFP, a marker of cardioblast-cardioblast contacts (Fig. 1f, Movie S3). We delineated individual cardioblasts and we approximated the position of each cell using its geometric centre (Fig. 1f’). We found that cardioblast velocities oscillated between positive and negative values. Consistent with our nuclear measurements, medially-directed cell steps were greater than laterally-directed steps in both amplitude (1.71±0.03-fold, *P* < 0.0001, Fig. 1g) and duration (1.76±0.05-fold, *P* < 0.0001, Fig. 1h). Together, these results indicate that, rather than monotonically advancing, cardioblasts repeatedly take forward and backward steps as they migrate.

**Figure 1.**
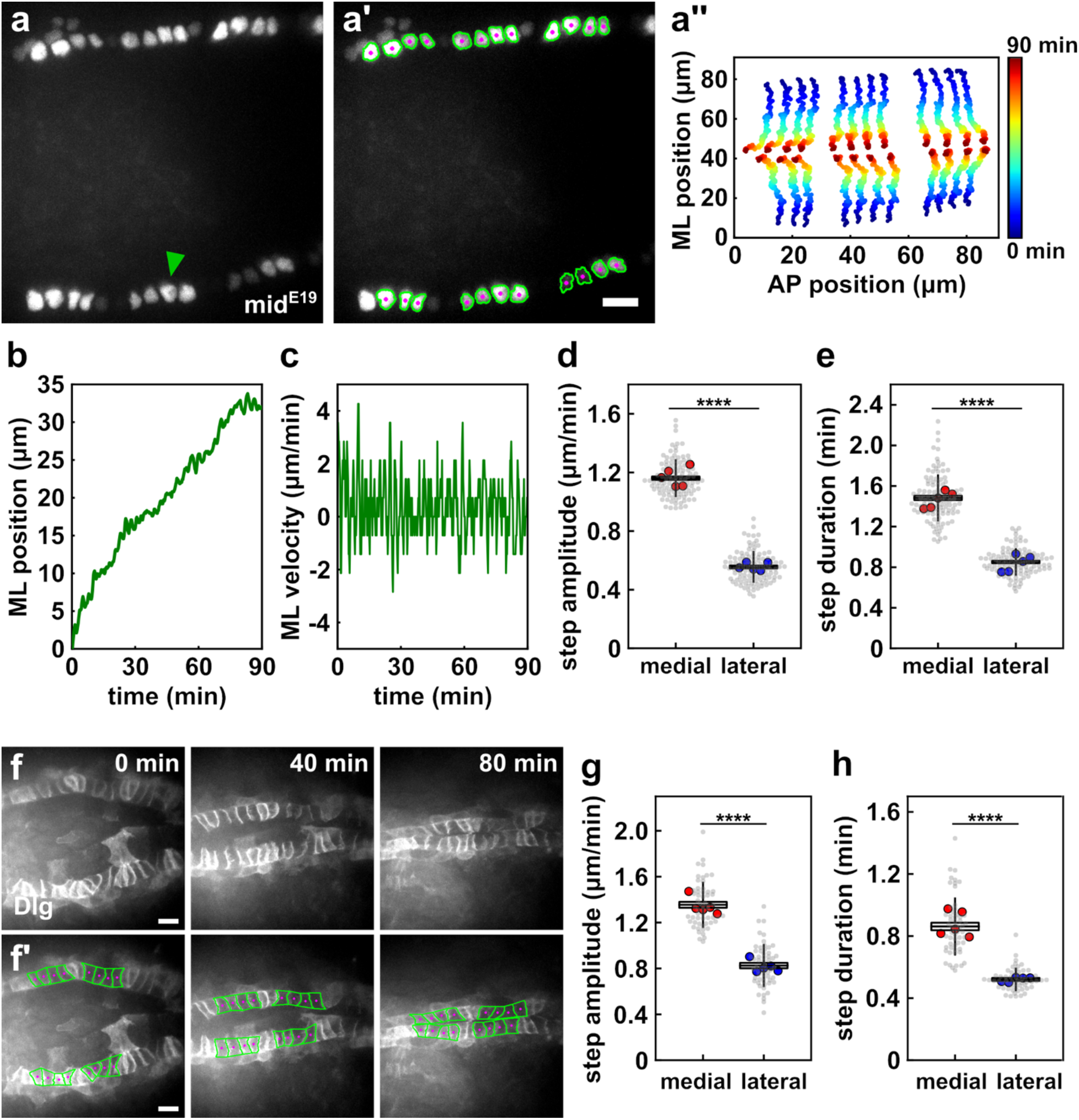
Cardioblast migration occurs in sequential forward and backward steps. **(a)** Cardioblast nuclei (a-a’) and their trajectories (a’’) in an embryo expressing midE19:GFP, a cardioblast-specific transcription factor. Green arrowhead (a) indicates cardioblast tracked in (b-c). Green indicates nuclear segmentation masks (a’). **(b-c)** Medial-lateral position (b) and velocity (c) of a cardioblast tracked over time. **(d-e, g-h)** Amplitude (d, g) and duration (e, h) of cardioblast forward and backward nuclear (d-e) or cellular (g-h) velocity steps. Grey points represent the mean for individual cells, and red/blue points represent embryonic means (*n* = 110 cells in 5 embryos in d-e, *n* = 58 cells in 5 embryos in g-h). Boxes indicate mean ± standard error of the mean (s.e.m.), error bars show the standard deviation (s.d.). **** *P* < 0.001, Wilcoxon signed-rank test. **(f)** Cardioblasts in an embryo expressing Dlg1:GFP (f), and corresponding cell segmentations (f’, green). Magenta labels cell centroids. (a, f) Anterior, left; medial, centre. Bar, 10 μm.

### Myosin waves oscillate between the leading and trailing edges of migrating cardioblasts

The molecular motor non-muscle myosin II generates mechanical forces critical for cell movement (Ridley et al., 2003; Franke et al., 2005). To determine if myosin contributes to the oscillatory migration of cardioblasts, we quantified myosin dynamics in embryos in which both cardioblasts and pericardial cells expressed GFP-tagged myosin. We found that myosin displayed a striking pattern of alternating localization between the leading and trailing edges of individual cardioblasts (Fig. 2a, Movie S4). The periodic pattern of myosin localization produced oscillatory waves that traversed the cells from front to back and *vice versa* (Fig. 2b). The period of myosin oscillations was 2.89±0.04 minutes, consistent with the periodicity of cardioblast movement. The period of myosin oscillation increased when contralateral neighbours made contact (2.67±0.07 minutes during migration, approximately one hour prior to contralateral cells making contact, *vs*. 3.16±0.05 minutes at the end of migration, *P* < 0.001, Fig. 2c-e), suggesting that myosin waves play a role in driving cardioblast movement. Cell movement in a 3-dimensional matrix often occurs in discrete steps. For example, in the *Drosophila* pupa, fat body cells migrate toward wound sites with peristaltic movement driven by retrograde actomyosin waves (Franz et al., 2018). Periodic actomyosin contractility is also necessary for the movement of fibroblasts within an elastic 3D matrix; repeated contractions at the leading edge of the cell pull the nucleus forward, resulting in periodic accelerations of the nucleus and dynamic changes in nuclear distance from the trailing edge of the cell (Petrie et al., 2014). Notably, our findings suggest that alternating forward and backward steps is also a viable form of *in vivo* cell migration.

**Figure 2.**
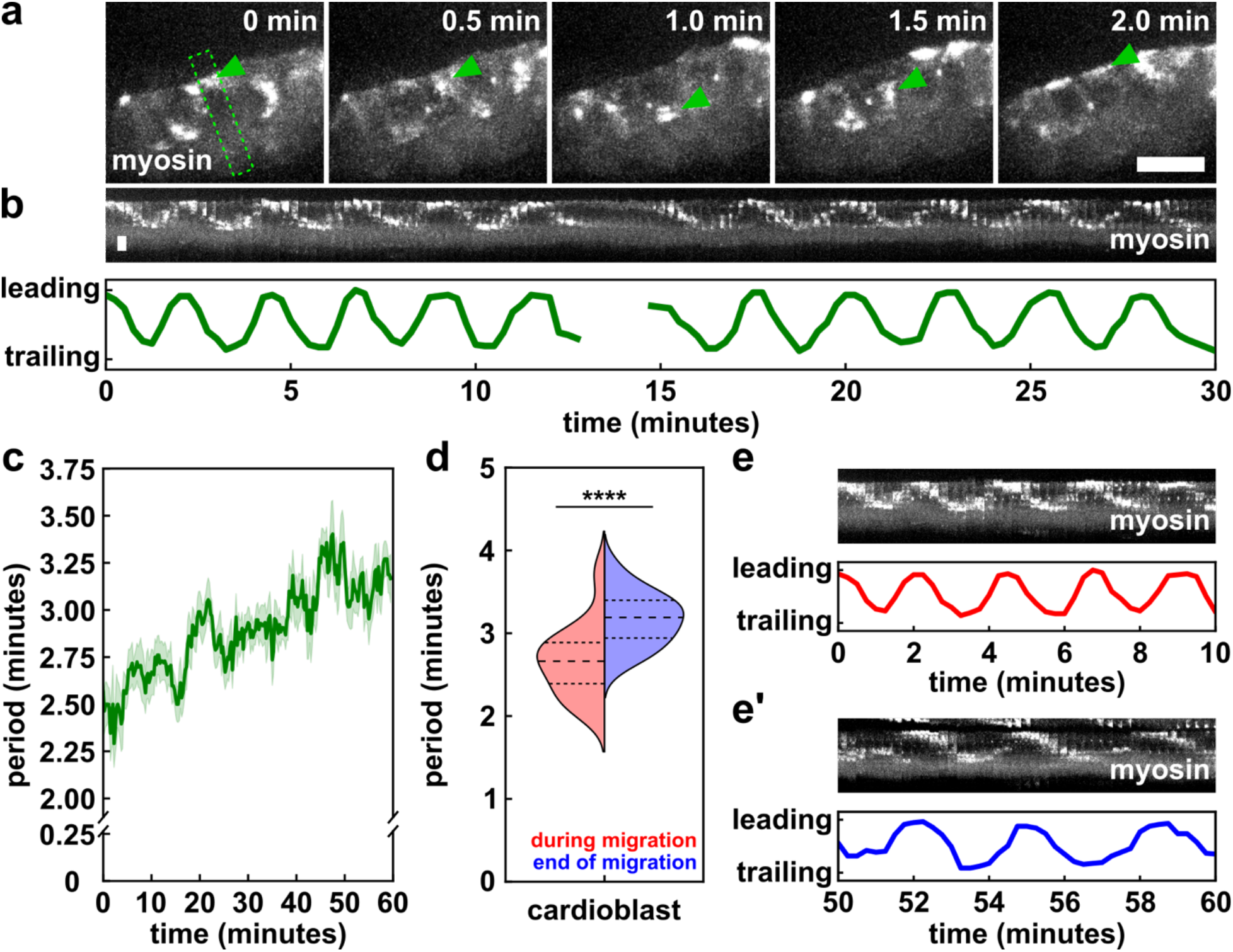
Cyclic myosin flows are associated with cardioblast migration. **(a)** Migrating cardioblasts expressing myosin:GFP. Green arrowheads track a myosin-rich structure moving from the leading to the trailing edge of a cell, and back to the leading edge. Green box indicates the region used to create the kymograph in (b). Anterior, left; medial, up. Bar, 10 μm. **(b)** Kymograph displaying myosin dynamics in the cardioblast shown in (a). (a-b) Time is with respect to the onset of the movie. **(c)** Myosin oscillation period during cell migration. Time is with respect to when contralateral partners make contact (at 60 min). Error bars, s.e.m. **(d)** Distribution of myosin oscillation period during cell migration (0-5 minutes in c, red, left), or when cardioblasts contact their contralateral partner (55-60 minutes in c, blue, right). Dashed lines indicate the first quartile, the median, and the third quartile. **(e-e’)** Representative kymographs displaying myosin dynamics during migration (e) and at the end of migration (e’) for the cardioblast shown in (a-b). Time is with respect to when contralateral partners make contact (at 60 min). (b, e) Medial, up. Bar, 15 s. (c-d) *n* = 46 cells in 5 embryos. **** *P* < 0.001, Wilcoxon signed-rank test.

### Myosin waves drive cell shape changes associated with cardioblast migration

To further investigate if myosin waves drive cardioblast migration, we measured the relationship between myosin flows and changes in position of cardioblast nuclei in embryos co-expressing GFP-tagged myosin and an mCherry-tagged nuclear localization sequence (Fig. S2a, Movie S5). We found that cyclic myosin flows were anti-correlated with the oscillations in the position of the cardioblast nucleus (Fig. S2b): when the myosin wave was at the leading edge of the cell, the nucleus shifted towards the trailing edge, and *vice versa*. Cross-correlation analysis revealed that changes in myosin localization preceded changes in nuclear position approximately by 20±5 seconds (Fig. S2c-d), further suggesting that fluctuating, myosin-based contractile forces drive cardioblast migration.

To determine if myosin waves cause cell shape changes in cardioblasts, we imaged heart development in embryos expressing Dlg1:GFP (Movie S3). We quantified the length of both leading and trailing edges, and we found that both ends of the cells underwent periodic contraction and relaxation (Fig. 3a-b). The period of contraction and relaxation was 3.1±0.1 minutes for the leading edge and 2.8±0.1 minutes for the trailing edge (Fig. 3c), consistent with the period of myosin fluctuation across the cell. Importantly, the contraction and relaxation of the leading and the trailing edge of a cardioblast were significantly anticorrelated (Fig. 3d-e). These results suggest that periodic cell shape changes may drive cardioblast movement.

**Figure 3.**
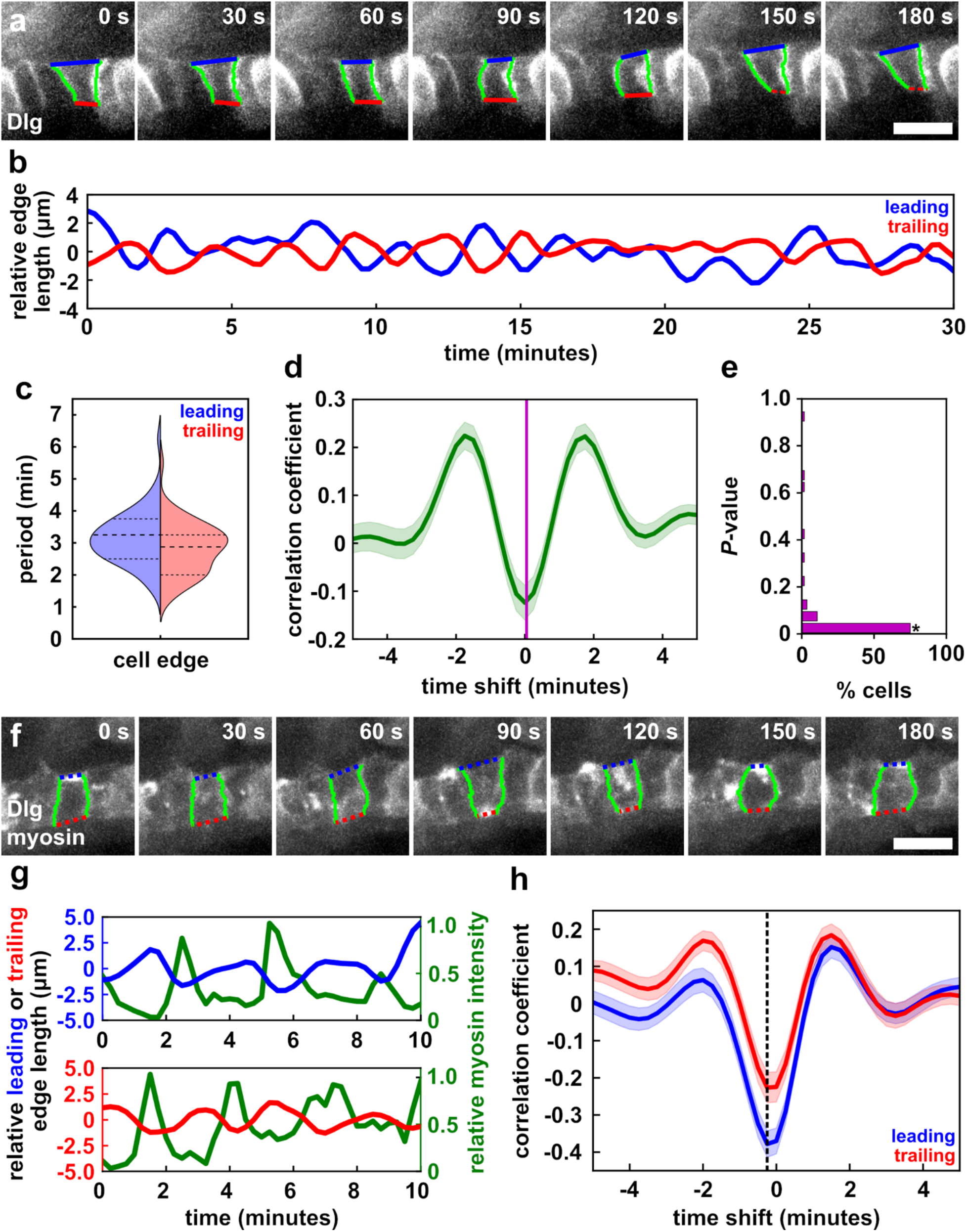
Myosin flows in cardioblasts are associated with periodic cell shape changes. **(a, f)** Cardioblasts expressing Dlg1:GFP (a), or Dlg1:GFP and myosin:GFP (f). Blue indicates the leading edge, red is the trailing edge, and green are the cardioblast-cardioblast contacts. **(b)** Length of leading (blue) and trailing (red) edges of the cardioblast shown in (a) relative to the mean length of the edge over time. **(c)** Period of length oscillation for the leading (blue) and trailing (red) edges of the cell. Dashed lines indicate the first quartile, the median, and the third quartile. **(d)** Cross-correlation measured when shifting the trailing edge length with respect to the leading edge length. Error bars, s.e.m. Magenta indicates the time shift corresponding to the minimum correlation. **(e)** *P-*value distribution for the anti-correlation between leading and trailing edge lengths. The bar labelled with an asterisk display *P* < 0.05. **(g)** Representative traces displaying length (blue and red) and myosin dynamics (green) for the leading (top) and trailing (bottom) edges of the cardioblast in (f). **(h)** Cross-correlation measured when shifting leading (blue) or trailing (red) edge length with respect to myosin position; dashed line represents the mean time shift corresponding to minimum correlation (*n* = 40 cells in 5 embryos). Error bars, s.e.m..

To determine if myosin was associated with cardioblast cell shape changes, we imaged cardiac development in embryos co-expressing GFP-tagged myosin and Dlg1 (Fig. 3f, Movie S6). We tracked the two markers based on their distinct localization patterns, with myosin predominantly excluded from contacts between cardioblasts (Fig. 2a), and Dlg1 almost exclusively present at the cell-cell contacts (Fig. 3a). We found that localization of myosin at the leading or trailing edges of cardioblasts correlated with the contraction of the corresponding edge (Figs. 3g-h and S3a-b). Specifically, cross-correlation analysis showed that myosin localization to the leading edge preceded the contraction of that edge by 9.4±1.5 seconds, and that myosin localization at the trailing edge preceded edge contraction by 8.3±1.5 seconds (Fig. S3a-b). Together, our data suggest that alternating, myosin-based contraction of the leading and trailing ends of individual cardioblasts drives cell shape changes that contribute to the peristaltic movement of cardiac progenitors during *Drosophila* heart development. The mechanisms that govern the alternative contraction and relaxation of leading and trailing cardioblast edges are unclear. Strain promotes myosin recruitment through the activation of mechanically-gated ion channels (Effler et al., 2006; Fernandez-Gonzalez et al., 2009; Zulueta-Coarasa and Fernandez-Gonzalez, 2018). Contraction of one cardioblast surface and the associated volume displacement could stretch the opposite cell surface, inducing myosin recruitment. The process would then repeat in the opposite cell surface, in a self-perpetuating cycle. The establishment of contacts with contralateral cardioblasts during lumen formation (Medioni et al., 2008; Santiago-Martínez et al., 2008) might limit myosin-induced deformation, thus interrupting the cycle.

### Cardioblasts are not strongly attached to the epidermis

Cardioblast migration occurs in parallel with dorsal closure, a process driven in part by the pulsatile contraction of amnioserosa cells (Solon et al., 2009; Blanchard et al., 2010; David et al., 2010). The amnioserosa is directly connected to the epidermis, and cardioblasts establish transient connections with epidermal cells (Rugendorff et al., 1994; Tepass and Hartenstein, 1994). Thus, we reasoned that pulsatile contraction in the amnioserosa could drive the oscillatory motion of cardioblasts. To determine whether external forces from the overlaying epidermis could contribute to the migration of cardioblasts, we examined the dynamics of cardioblast movement relative to the epidermis. We imaged embryos co-expressing the adherens junction marker E-cadherin:mKate as an epidermal cell outline marker, and mid^E19^:GFP to label the cardioblast nuclei. To induce rapid displacements in the epidermis, we used laser ablation to sever the interface between the epidermis and the amnioserosa, a site of increased tension (Hutson, Kiehart, 2000) (Fig. S4a). Laser ablation induced recoil movements in the epidermis, quantifiable as 2-fold greater lateral velocity of the tricellular junctions around leading edge epidermal cells immediately after ablation as compared to sham-irradiated controls (*P* < 0.0001, Fig. S4b). In contrast, the cardioblast nuclei closest to the epidermal cells impacted by the ablation did not display a change to their lateral velocity when compared to sham-irradiated controls (Fig. S4c). Our findings are consistent with previous results showing that cardioblasts move faster than leading edge epidermal cells, and that in dorsal closure mutants cardioblasts overtake the leading edge of the epidermis (MacMullin and Jacobs, 2006; Haack et al., 2014). Together, these results indicate that the connections between cardioblasts and epidermal cells are not strong, suggesting that cardioblasts do not move anchored to the overlaying epidermis.

### Modelling predicts that a stiff trailing edge boundary biases the direction of cardioblast movement

To investigate how the alternative and periodic contraction of the leading and trailing edges could drive cardioblast movement in the absence of external forces, we developed a mathematical model of cardioblast migration (see Methods). We used the vertex modelling framework, in which cells are represented by their corners, and differential equations calculate the movement of each cell corner over time (Nagai and Honda, 2001; Yu and Fernandez-Gonzalez, 2017). We initially developed a simple model in which cells preserved their volumes and the leading and trailing edges of the cells contracted alternatively, with periods corresponding to those measured *in vivo* (Fig. 2c-e). Under these conditions, we found that the cell centroid position fluctuated medially and laterally, but the cell did not make any net progress in its migration (Fig. 4a and Movie S7, top). Thus, we speculated that some form of asymmetry was necessary to bias the movement of the cell medially.

**Figure 4.**
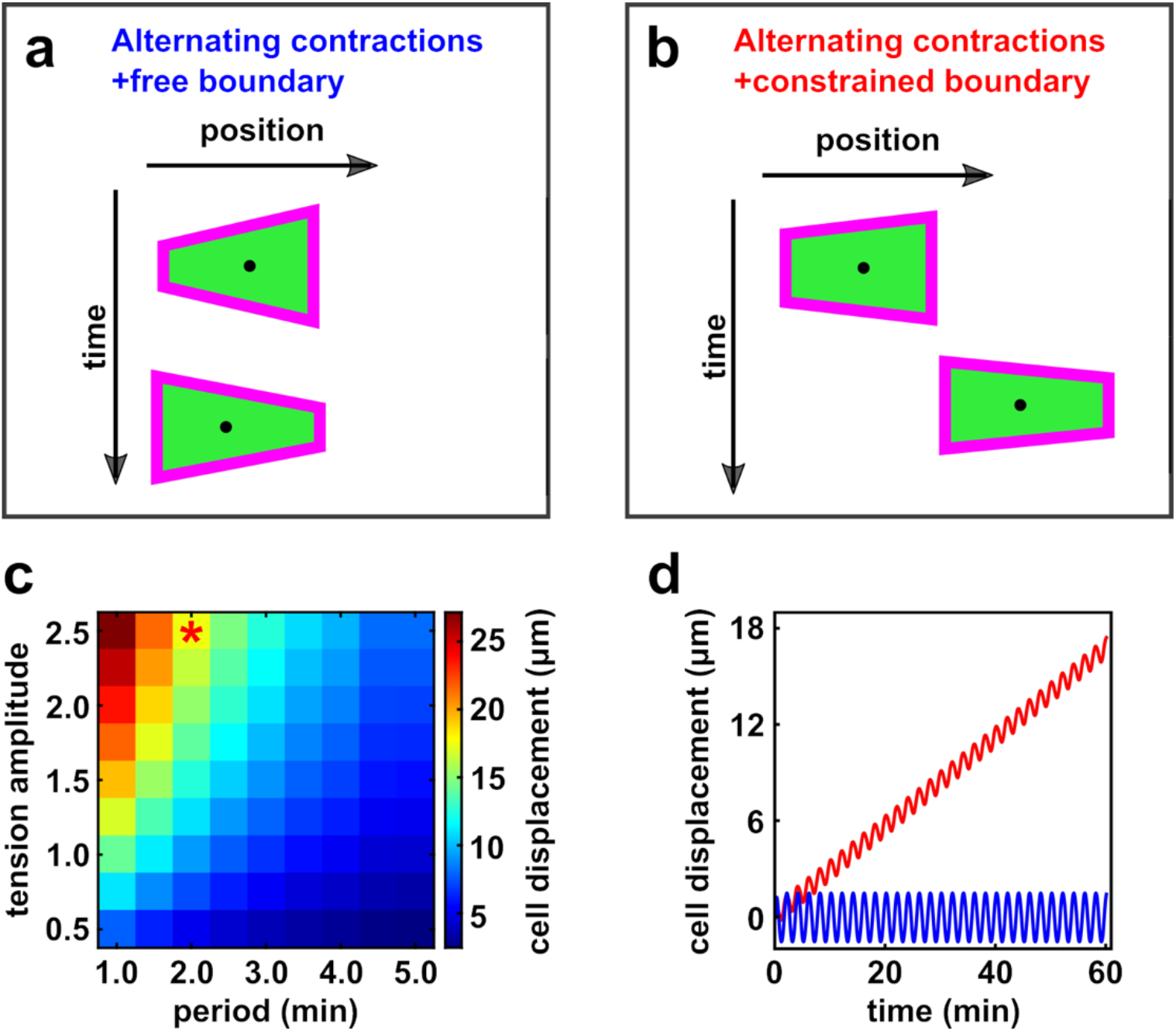
A mechanical asymmetry can bias the direction of peristaltic cell movements. (**a-b)** Simulations of cardioblast migration with free leading edges, and free (a) or constrained (b) trailing edges. **(c)** Cell displacement (net forward movement of the cell centre) within 60 minutes as a function of tension amplitude (*γ*_*0*_) and oscillation period (*T*). Asterisk highlights the simulation values used in (d). **(d)** Cell displacement over time for free (blue) and constrained (red) trailing edges, for *γ*_*0*_ = 2.5 and *Τ* = 2 min.

Stiff borders between cell populations can limit movements and prevent cell mixing (Umetsu and Dahmann, 2010). To investigate the role that mechanical asymmetries may play in cardioblast migration, we introduced a boundary at the trailing edge of the cell in our simulations. The presence of a stiff boundary, coupled with the periodic and alternative contraction of leading and trailing edges, were sufficient to produce oscillatory, net medial movement of the cell (Fig. 4b-d and Movie S7, bottom), reminiscent of our *in vivo* observations (Fig. 1b). Thus, mathematical modelling predicts that a boundary at the trailing edge may bias the direction of cardioblast migration medially.

To test the prediction that the trailing edge of cardioblasts is stiffer than the leading edge, we quantified the deformability of cardioblast leading and trailing edges *in vivo* by measuring their amplitude of contraction and expansion (Fig. 1f, Movie S3). We found that during cardioblast migration, the deformation of the leading edge was significantly greater than the deformation of the trailing edge (Fig. S5a-b). Contractions of the leading edge were on average 22±5% greater *(P* < 0.001, Fig. S5a), and expansions of the leading edge were 24±5% greater *(P* < 0.001, Fig. S5b). Our results suggest that the trailing edge of the cardioblasts is more resistant to deformation than the leading edge.

To investigate if the reduced deformability of the trailing edge is linked to a limitation in the lateral movement of the cardioblasts, we quantified the medial and lateral displacement of the vertices that delimit the leading and trailing edges (Fig. S5c, Movie S3). We found that the medial movement of leading edge vertices was 53±3% greater than their lateral movement (*P* < 0.0001, Fig. S5c). Similarly, the medial movement of trailing edge vertices was 69±4% greater than their lateral movement (*P* < 0.0001, Fig. S5c). Together, these results suggest that the backward movement of cardioblasts is restricted by a stiff boundary at their trailing edge.

### A supracellular actin cable separates cardioblasts and pericardial cells and limits backward cardioblast movement

Actin networks can change the material properties of cell surfaces (Blanchoin et al., 2014). To determine what is the molecular mechanism that reduces the deformability of the trailing edge of cardioblasts, we examined actin dynamics. We used embryos expressing GFP:MoesinABD in cardiac progenitors as an F-actin reporter (Haack et al., 2014). Strikingly, we found that a supracellular actin cable formed at the trailing edge of the cardioblasts, at the interface with the pericardial cells (Fig. 5a-a’, Movie S8), possibly creating a boundary between the two cell types. Actin levels were 28±3% greater at the trailing edge of cardioblasts than at the leading edge (*P* < 0.005, Fig. 5b), suggesting that the accumulation of actin might stiffen the trailing edge of the cardioblasts, establishing a boundary that limits lateral movement.

**Figure 5.**
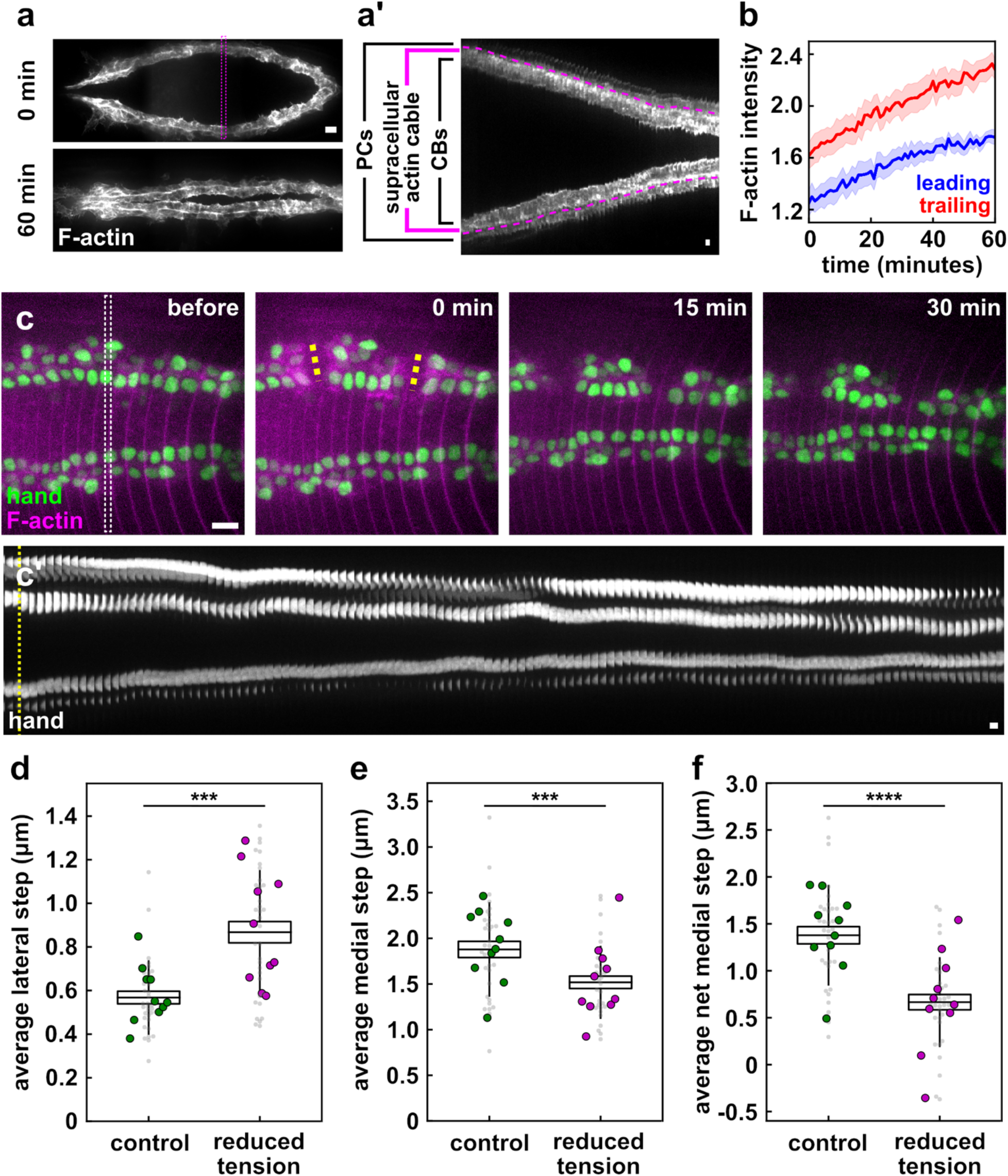
A supracellular actin cable promotes medial cardioblast migration. **(a-a’)** Migrating cardioblasts expressing GFP:MoesinABD (a marker of filamentous actin). Pink box represents the region used to create the kymograph in (a’). **(b)** F-actin fluorescence at the leading (blue) or trailing (red) edge of cardioblasts (*n* = 12 cells in 6 embryos), error bars, s.e.m. **(c)** Embryo expressing Hand:GFP (green) and mCherry:MoesinABD (magenta), before (left) and after (right panels) ablation of the actin cable. Dotted yellow lines indicate ablation sites. White box shows the region used to create the kymograph in (b’), where a dotted yellow line indicates the time of ablation. **(d-f)** Mean lateral step (d), medial step (e), and net medial displacement per medial-lateral cycle (f) in control (green) and ablated (magenta) cells. Boxes indicate mean ± s.e.m., error bars show the s.d.. Grey points represent the mean for individual cells, and green/magenta points represent embryonic means (*n* = 34 cardioblast pairs in 10 embryos). *** *P* < 0.005, **** *P* < 0.001, Wilcoxon signed-rank test. (a-b) Anterior, left; medial, centre. Bar, 10 μm in (a, c), 1 minute in (a’), and 15 seconds in (c’).

To determine whether the supracellular actin cable limits the lateral movement of the cardioblasts, we severed the actin cable in embryos expressing Hand:GFP to label cardioblast and pericardial cell nuclei (Han and Olson, 2005), and mCherry:MoesinABD to label the actin cable (Millard and Martin, 2008) (Fig. 5c, Movie S9). We cut two lines perpendicular and across the cable to reduce the tension that the cable sustained (Figure 5c, yellow dashed lines). We found that after reducing cable tension, the oscillation dynamics of the cardioblasts were disrupted (Fig. 5c’, Movie S9). Reducing tension at the supracellular actin cable resulted in a 64±25% increase in the amplitude of the lateral steps of the cardioblasts with respect to the contralateral, non-ablated cardioblasts (Fig. 5d, *P* < 0.0001), and a 10±8% decrease in the amplitude of the medial steps (Fig. 5e, *P* < 0.005). The changes in oscillatory dynamics had consequences for the migration of cardioblasts: disrupting the actin cable reduced the net medial movement per cycle of medial and lateral steps by 43±17% (Fig. 5f, *P* < 0.001). These results indicate that the supracellular actin cable at the interface between cardioblasts and pericardial cells restricts lateral cardioblast movement, biasing cardioblast migration medially.

Our findings show that coupling a stiff boundary at the trailing edge of a cell with periodic shape changes can drive directional cell movements *in vivo*. In cardioblasts, myosin waves generate the energy to move, and the actin cable at the trailing edge steers the movement. The oscillatory migration of cardioblasts is reminiscent of the amoeboid movement of slime molds (Zhang et al., 2017) or the periodic migration of single human cells depleted of the actin-associated protein zyxin in 3D culture (Fraley et al., 2012). However, cardioblasts move collectively, maintaining stable contacts with their anterior and posterior neighbours (Haag et al., 1999; Wang et al., 2005). The contributions of individual cell shape changes and how these changes are coordinated to facilitate the collective migration of cardiac progenitors remains an open question.

## Materials and methods

### Fly stocks

We used the following markers for live imaging: E-cadherin:mKate2×3 (Pinheiro et al., 2017), *hand-hand*:*GFP* (Han and Olson, 2005), *mid*-*mid*^E19^*:GFP* (Jin et al., 2013), *hand-GFP:moesinABD* (Haack et al., 2014), *UAS-dlg1:GFP* (Koh et al., 1999), *UAS-mCherry:moesinABD* (Millard and Martin, 2008), *UAS-mCherry:nls* (Bloomington *Drosophila* Stock Center #38424), and *UAS-*s*qh:GFP* (gift of E. Caussinus). UAS constructs were driven with *hand*-Gal4 (Bloomington *Drosophila* Stock Center #48396), or *tinCΔ4*-Gal4 (Lo and Frasch, 2001).

### Time-lapse imaging

Stage 14 embryos were dechorionated in 50% bleach for 2 minutes, rinsed, glued dorsal side down to a glass coverslip using heptane glue, and mounted in a 1:1 mix of halocarbon oil 27 and 700 (Sigma-Aldrich, St. Louis, MO)(Scepanovic et al., 2021). Embryos were imaged using a Revolution XD spinning disk confocal microscope equipped with an iXon Ultra 897 camera (Andor, Belfast, UK), and using a 60x oil immersion lens (Olympus, Shinjuku, Japan; NA 1.35). Sixteen-bit Z-stacks were acquired at 0.75 μm steps every 15-30 s (21-27 slices per stack).

Maximum intensity projections were used for analysis in the heart. In the epidermis, Z-stacks were projected using a local Z projector to correct for the curvature of the embryo (Herbert et al., 2021). To image the dynamics along the entire length of the heart, 3 overlapping positions along the embryo were imaged as above and maximum intensity projections were stitched using a feathering blending algorithm, where the weighting coefficients at the stitching seam between two input images were calculated based on the distances of each input from the seam (Uyttendaele et al., 2001).

### Laser ablation

Laser cuts were created with a pulsed Micropoint N2 laser (Andor) tuned to 365 nm. The laser delivers 120 μJ pulses of 2-6 ns each. For ablation of the supracellular actin cable at the interface between cardioblasts and pericardial cells, 10 consecutive pulses were delivered at discrete points ~ 2 μm apart along two 14 μm lines perpendicular to the cable. For cuts along the leading edge of the epidermis, 10 consecutive pulses were delivered at discrete points ~ 2 μm apart along a 14 μm line at the boundary between the amnioserosa the epidermis. For controls, the laser was fully attenuated using a neutral density filter.

### Cell segmentation and image analysis

Image analysis was performed using our open-source image analysis platforms, PyJAMAS (Fernandez-Gonzalez et al., 2021) and SIESTA (Fernandez-Gonzalez and Zallen, 2011). To automatically detect the position of cardioblast nuclei from microscopy movies, we used a support vector machine, a supervised machine-learning algorithm (Wang and Fernandez-Gonzalez, 2017), to distinguish cell nuclei from the background. We trained a linear support vector machine using a collection of 46×46 pixel images containing 91 images of cardioblast nuclei (positive training set) and 101 background images (negative training set), with 512 features per image. Image features were based on the histogram of gradients, which describes the shape of an object using the distribution of local image gradient orientations (Fernandez-Gonzalez et al., 2021). The support vector machine identified the linear boundary that separated nuclear from background images in feature space. The linear boundary was then applied to the feature set calculated for 46×46 pixel regions of interest obtained from new images to determine if the region of interest corresponded to a nucleus or to the background. Regions of interest were sequentially shifted by 1 pixel in the X or Y direction to fully scan the new images. Each image was assigned a probability of containing a nucleus obtained using 5-fold cross-validation and Platt scaling (Platt, 1999). Non-optimal and overlapping detections were eliminated using non-maximum suppression by setting the maximum intersection over union to 0.3, and the minimum probability of containing a nucleus to 0.95. Each detected nucleus was automatically segmented using the watershed algorithm, a region growing method (Beucher, 1992). Briefly, a seed point was set in the brightest pixel within each of the windows containing a nucleus (nuclear seed), and a second seed point was set in the dimmest corner of each window (background seed). Seed points were grown on the gradient transform of the image window, simulating a flooding process in which pixels were assigned to seeds based on their increasing pixel value, until the two grown seeds met at the edge of the nucleus. We used the geometric centre of each segmented nucleus to represent nuclear position. Nuclei were tracked across time points by matching the centroid of the closest nuclei in consecutive images. The migration dynamics of cardioblasts was quantified using the medial-lateral component of cell velocities.

To segment cardioblasts expressing Dlg1:GFP, we used the LiveWire algorithm in PyJAMAS, an interactive method based on Dijkstra’s minimal cost path algorithm (Dijkstra, 1959) to find the brightest path between to pixels. The leading and trailing edges of the cells, where Dlg1:GFP does not localize were closed using a straight line.

The oscillation period was quantified as the dominant frequency of the signal autocorrelation, calculated using a fast Fourier transform. For intracellular myosin localization, period was measured for 10-minute segments of uninterrupted oscillations extracted from kymographs of myosin localization. Mean myosin fluorescence at the leading and trailing edges of the cell was measured from maximum intensity projections, and background-subtracted using the most frequent pixel value (the mode) of the image in each time point. Intensity values were corrected for photobleaching by dividing by the mean intensity of the image in each time point.

### Mathematical modeling of cardioblast migration

To investigate how the periodic cell shape changes experienced by cardioblast could drive cell movement, we developed a single cell vertex model of cardioblast migration in two dimensions. The energy, *E*, of the cell can be defined as:

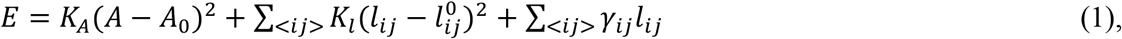

where the first term represents the cell volume incompressibility, the second term represents the spring restoring energy of the cell edges, and the last term denotes myosin-based contractility. In Eq. 1, *K*_*A*_ and *K*_*l*_ are the area and linear spring constants; *A* and *A*_0_ are the actual and preferred cell areas, respectively; and *l*_*ij*_ and 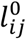 are the distance between vertices *i* and *j* and the rest length of the edge connecting vertices *i* and *j*, respectively (we assumed that each cell edge was a spring). *γ*_*ij*_ represents the myosin-generated line tension on the edge connecting vertices *i* and *j*.

To model the periodic myosin waves, we assigned a sinusoidal and anti-correlated *γ*_*i,j*_ for the leading and trailing edges:

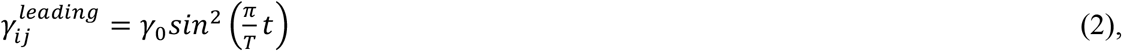

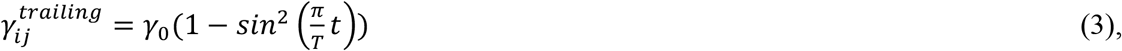

where *γ*_0_, the tension amplitude, is proportional to the total myosin flowing in the cell and represents the maximum magnitude of the myosin oscillations; and *T* is the period of myosin oscillations at the leading and trailing edges. We set *γ*_*ij*_ = 0 for the anterior and posterior edges of the cell.

The evolution of the cell is guided by an energy minimization process. At each time step of the simulations, we updated the position of the vertices using the forward Euler method:

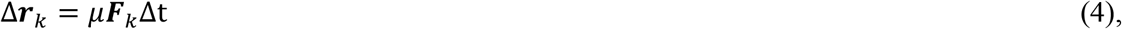

where ***r***_*k*_ is the position vector of vertex *k*, ***F***_*k*_ = −∇_*k*_*E* is the force on vertex *k*, *μ* is the inverse friction coefficient, and Δ*t* is the integration time step. We set *μ* = 1 for all the simulations. The integration time step was Δ*t* = 0.01*τ*, where *τ* = 1/(*μK*_*A*_*A*_*0*_) is the natural time unit of the simulations.

We started our simulations with a rectangular cell, with initial leading and trailing edge lengths of 0.8 *l*, and anterior and posterior edge lengths of 1.25 *l*, where *l* is the natural length unit of the simulations, 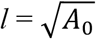. The same values were assigned as rest lengths. We set the preferred area of the cell to *A*_0_ = *l* ^2^. We assumed that the cell size *A* did not change over time, and we selected a relatively high *K*_*A*_ value (*K*_*A*_ = 10) to ensure that the cell area size remained close to its target value. We set the linear spring constant to be the same value for all edges, *K*_*l*_ = 1 (Fig. 4a and Movie S7, top), or to *K*_*l*_ = 5 for the trailing edge and *K*_*l*_ = 1 for all other edges to simulate a mechanical asymmetry (Fig. 4b and Movie S7, bottom). *γ*_*0*_ and *T* were the two free parameters in our simulations that we explored using a parameter sweep (Fig. 4c). In addition, to simulate the presence of a boundary at the trailing edge, we restricted the medial-lateral movement of trailing edge vertices to the medial direction.

To convert the natural unit length of the simulations to micrometres, we measured the length of anterior and posterior cardioblast edges. The average anterior and posterior edge length was 9.9±0.8 μm (*n* = 56 cells). As the rest length for the anterior and posterior edges in the simulations was 1.25 *l*, we approximated *l* ~ 8 μm. We run the simulations for 6000 steps (60 *τ*) and assigned a relationship between the natural time unit of the simulations, *τ*, and minutes as *τ* = 1 min, as the net medial movement of the cell at 60 minutes *in vivo* (Fig. 1b) was comparable to that at 6000 steps *in silico* (Fig. 4d).

### Statistical analysis

Sample means were compared using non-parametric Mann-Whitney tests, or Wilcoxon signed-rank tests for paired data. For time series, error bars indicated the standard error of the mean (s.e.m.). For box plots, error bars show the s.d., boxes indicate the s.e.m., and the mean is indicated as a line inside of the box.

## Acknowledgments

We wish to acknowledge this land on which the University of Toronto operates. For thousands of years it has been the traditional land of the Huron-Wendat, the Seneca, and the Mississaugas of the Credit. Today, this meeting place is still the home to many Indigenous people from across Turtle Island and we are grateful to have the opportunity to work on this land. We are grateful to Andrew Renault, Emmanuel Caussinus, Zhe Han, Eurico Morais-de-Sá, and Georg Vogler for reagents. We thank Ana Maria do Carmo, Tony Harris, Michelle Ly, Gordana Scepanovic, Ian Scott and Teresa Zulueta-Coarasa for comments on the manuscript. This work was funded by the Natural Sciences and Engineering Research Council of Canada (418438-13), the Canada Foundation for Innovation (30279), the Translational Biology and Engineering Program of the Ted Rogers Centre for Heart Research, and the Canadian Institutes of Health Research (156279). N.B. was supported by an Ontario Graduate Scholarship. C.M. was supported by a Canada Graduate Scholarship-Doctoral from the Natural Sciences and Engineering Research Council of Canada. R.F.-G. is the Canada Research Chair in Quantitative Cell Biology and Morphogenesis.

## Supplementary figure legends

**Supplementary Figure 1.**
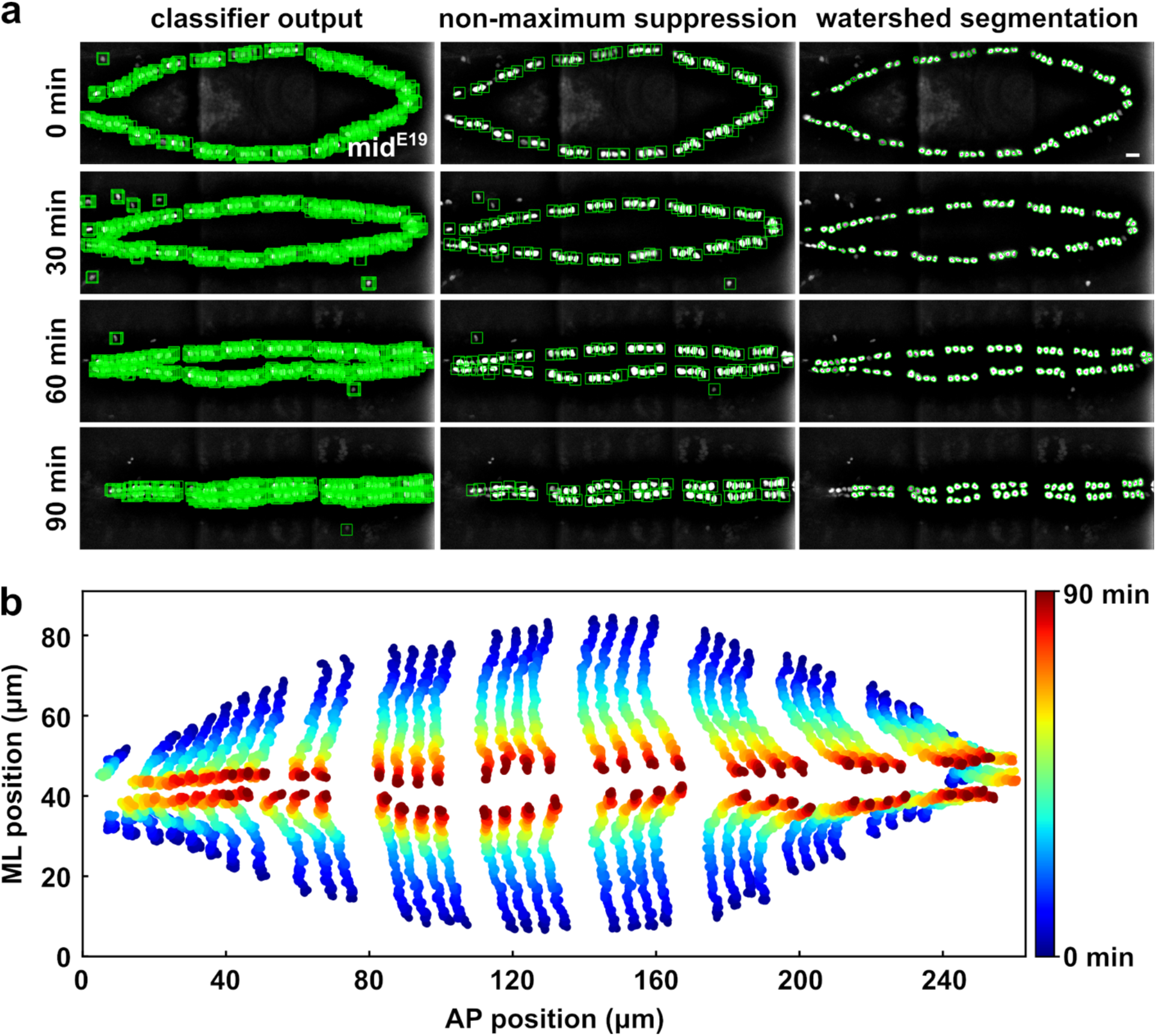
Cardioblast segmentation and tracking using a support vector machine. **(a)** Detection and segmentation of cardioblast nuclei in an embryo expressing mid^E19^:GFP. Green boxes indicate nuclei detected by a support vector machine trained to detect cardioblast nuclei, before (left) and after (middle) non-maximum suppression; green outlines delineate the nuclei (right). Magenta labels nuclear centroids. Bar, 10 μm. **(b)** Cardioblast trajectories for the embryo in (a). (a-b) Anterior, left; medial, centre.

**Supplementary Figure 2.**
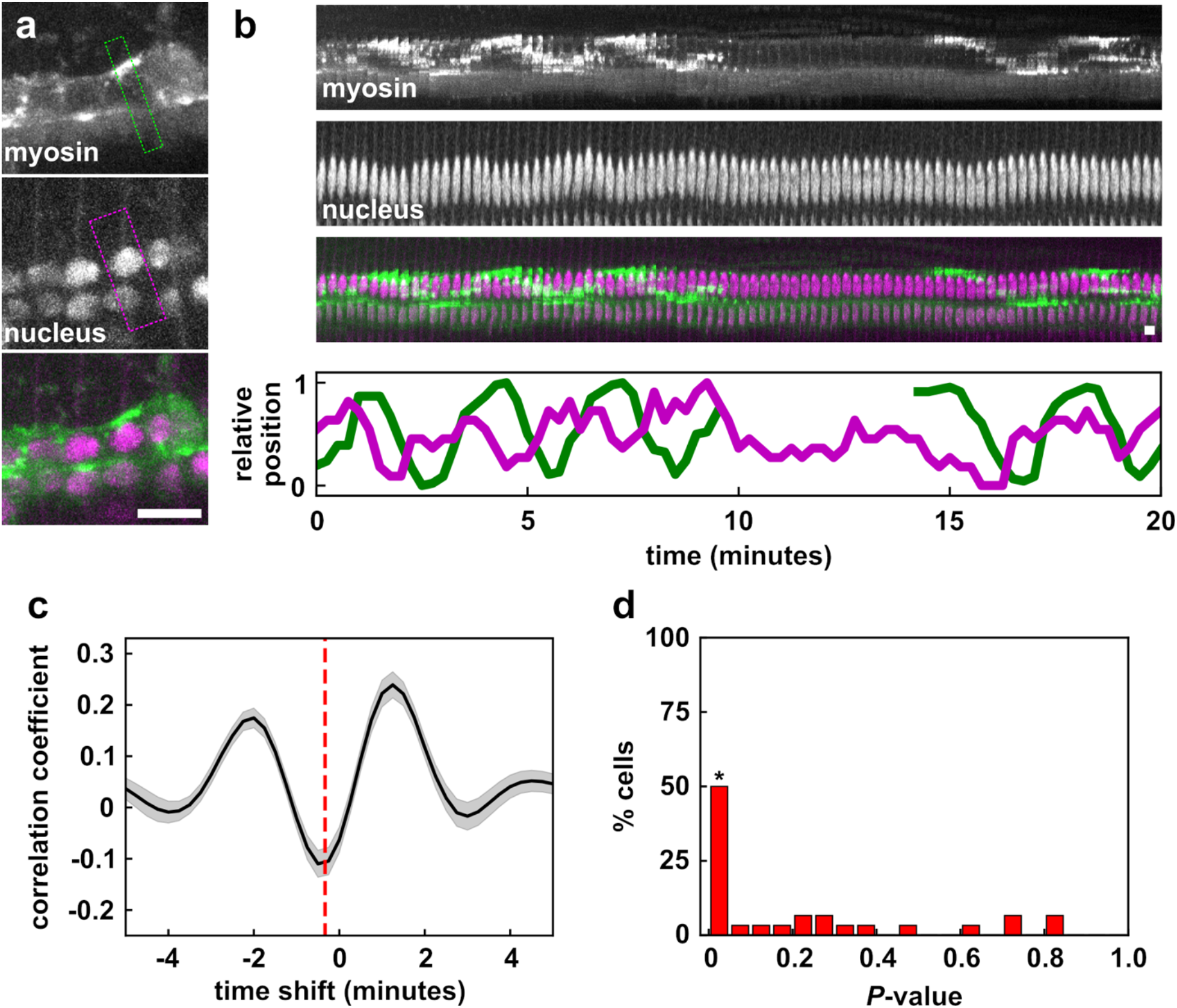
Myosin flows precede the movement of cardioblast nuclei. **(a)** Migrating cardioblasts expressing myosin:GFP (top, green in overlay) and an mCherry-tagged nuclear localization sequence (middle, magenta in overlay). Dashed boxes indicate the region used to create the kymographs in (b). Anterior, left; medial, up. Bar, 10 μm. **(b)** Kymographs displaying myosin (top, green in overlay) and nuclear position (center, scaled vertically by a factor of 2 to reveal oscillations, magenta in overlay), and traces displaying myosin (green) and nuclear (magenta) dynamics (bottom). **(c)** Cross-correlation measured when shifting nucleus position with respect to myosin. Error bars, s.e.m.. Dashed line indicates the time shift corresponding to the minimum correlation. **(d)** *P-*value distribution for minimum correlations between myosin and cardioblast nucleus position. Bars labelled with an asterisk display *P* < 0.05. (c-d) *n* = 30 cells in 5 embryos.

**Supplementary Figure 3.**
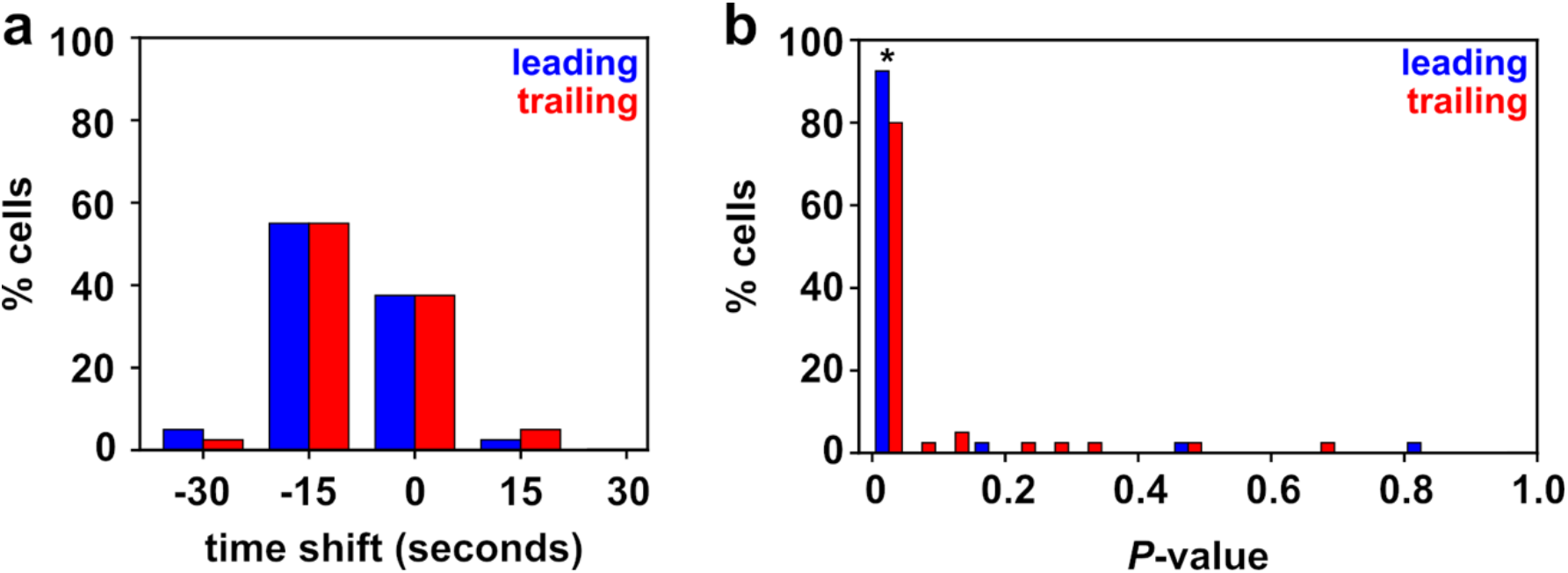
Myosin localization is anti-correlated with edge length. **(a)** Distribution of time shifts resulting in minimum cross-correlation when shifting leading (blue) or trailing (red) edge length relative to myosin fluorescence at the respective edge. **(b)** *P*-value distribution for minimum correlations between myosin fluorescence and edge length at the leading (blue) or trailing (red) ends of cardioblasts. The bar labelled with an asterisk displays *P* < 0.05. (a-b) *n* = 40 cells in 5 embryos.

**Supplementary Figure 4.**
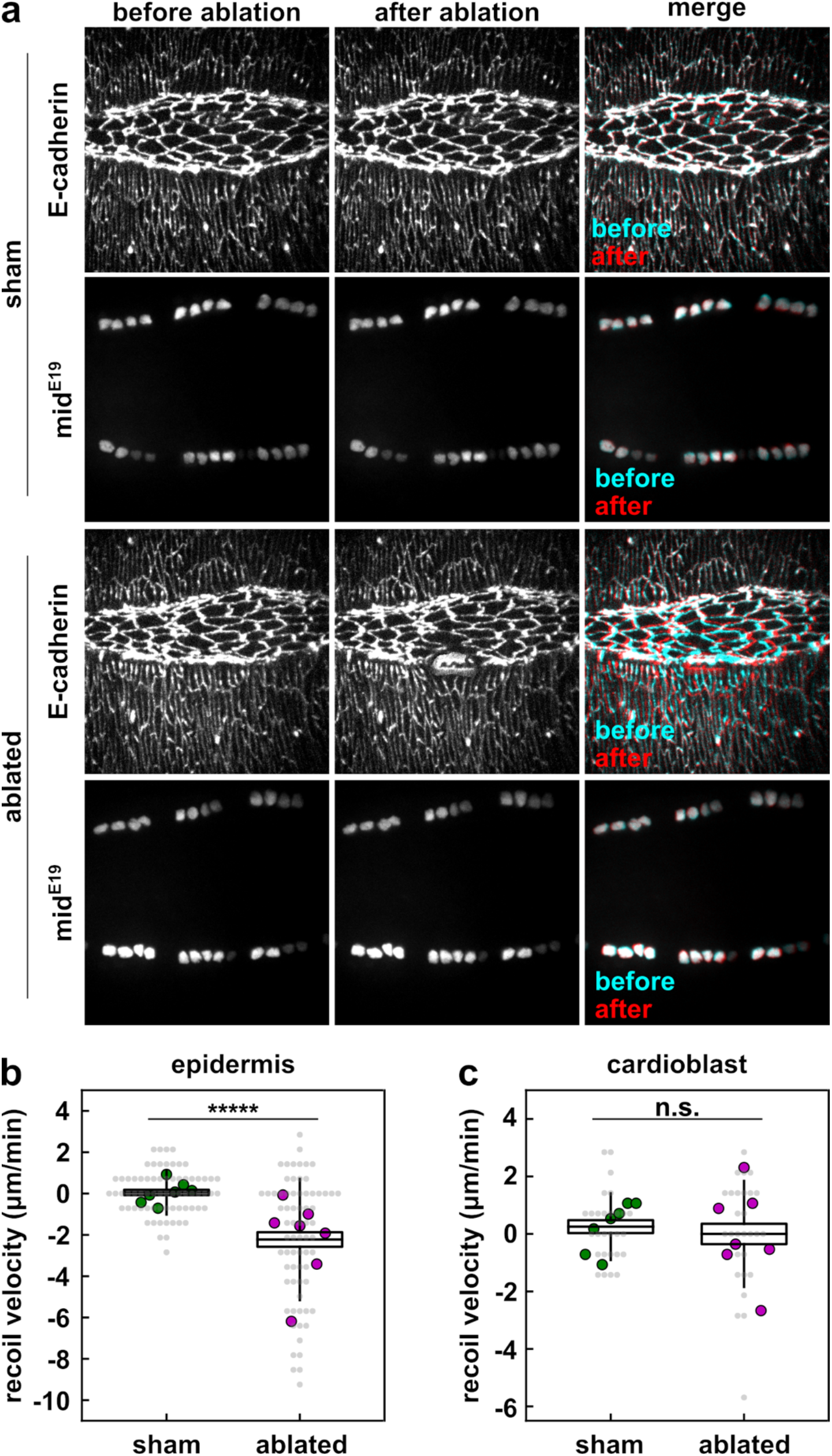
Cardioblasts move independently from the epidermis. **(a)** Embryos co-expressing E-cadherin:mKate (top) and mid^E19^:GFP (bottom), before (left, cyan in merge) and 15 s after (centre, red in merge) laser ablation at the epidermal leading edge. Anterior, left. **(b-c)** Instantaneous medial-lateral retraction velocity of tricellular junctions of leading edge epidermal cells (b, *n* = 70 junctions in 7 embryos for sham, *n* = 70 in 7 embryos for ablated) or cardioblasts closest to the site of ablation (c, *n* = 28 cardioblasts in 7 sham embryos, *n* = 28 in 7 ablated embryos). Grey points represent the mean for individual tricellular vertices (epidermis) or individual cells (cardioblasts), and magenta/green points represent embryonic means. Boxes indicate mean ± s.e.m., error bars show the s.d.. ***** *P* < 0.0001, n.s. not significant, Mann-Whitney test.

**Supplementary Figure 5.**
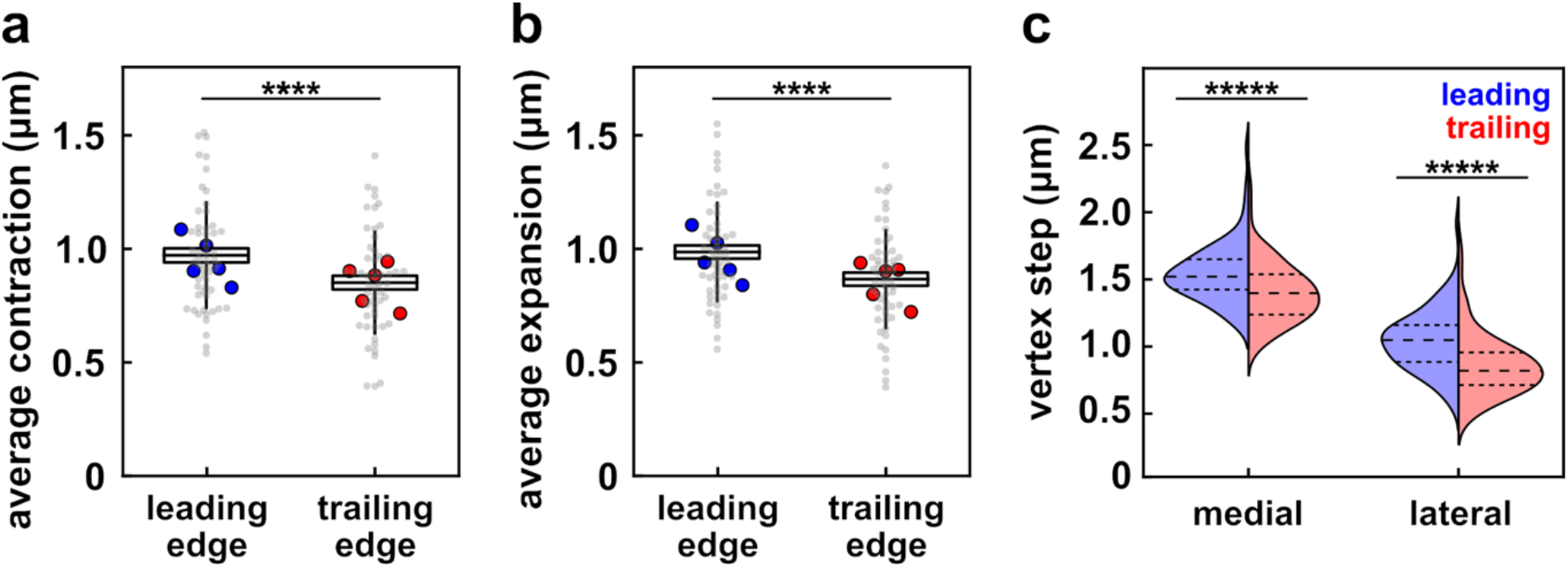
The trailing edge of cardioblasts resists deformation. **(a-b)** Average contraction (a) and expansion (b) of the leading (blue) and trailing (red) edges of cardioblasts. **(c)** Forward and backward displacement per step of vertices at the leading (blue) and trailing (red) edges of cardioblasts. Dashed lines indicate the first quartile, the median, and the third quartile. (a-b) Grey points represent the mean for individual cells, and red/blue points represent embryonic means. Boxes indicate mean ± s.e.m., error bars show the s.d.. (a-c) *n* = 56 cells in 5 embryos. **** *P* < 0.001, ***** *P* < 0.0001, Wilcoxon signed-rank test.

## Supplementary movie legends

**Movie S1. Cardioblasts migration occurs in steps.** Heart development in an embryo expressing the cardioblast-specific nuclear marker *mid*^E19^:GFP. The central region of the heart was imaged every 15 s. Anterior, left; medial, centre. Green indicates nuclear outlines.

**Movie S2. Tracking cardioblast migration using a support vector machine.** Heart development in an embryo expressing the cardioblast-specific nuclear marker *mid*^E19^:GFP. Three stacks covering the length of the heart were acquired every 15 s and stitched together. Cardioblast nuclei were detected using a support vector machine (top panel), non-maximum suppression was used to automatically select one box per nucleus (second panel), and nuclei were segmented using watershed algorithm (third panel). Migration trajectories (bottom panel) were reconstructed as time projections of the position of the nuclear centroid (third panel, magenta).

**Movie S3. Cardioblasts undergo periodic shape changes as they migrate.** Heart development in an embryo expressing *UAS-dlg1:GFP* driven by *tinCΔ4-Gal4*. The central region of the heart was imaged every 15 s. Anterior, left; medial, centre.

**Movie S4**. **Myosin flows across cardioblasts as they migrate.** Heart development in an embryo expressing *UAS-sqh:GFP* driven by *hand-Gal4*. The posterior region of the heart was imaged every 15 s. Anterior, left; medial, centre. Green arrowhead indicates the cell shown in Fig. 2a.

**Movie S5. Myosin flows are associated with nuclear oscillations.** Heart development in an embryo expressing *UAS-sqh:GFP* (green) and *UAS-mCherry:nls* driven by *hand-Gal4*. The central region of the heart was imaged every 15 s. Anterior, left; medial, centre.

**Movie S6. Myosin flows are associated with periodic cardioblast shape changes.** Heart development in an embryo expressing *UAS-dlg1:GFP* and *UAS-sqh:GFP* driven by *tinCΔ4-Gal4* and *hand-Gal4*. The central region of the heart was imaged every 15 s. Anterior, left; medial, centre.

**Movie S7. Modelling predicts that a stiff boundary at the trailing edge and periodic cell shape changes propel cardioblasts forward.** Vertex model simulations of a symmetric cardioblast that maintains a constant area and undergoes periodic and alternative contraction and relaxation of leading and trailing edges (top), and after the addition of a stiff boundary at the trailing edge (bottom), for *γ*_*0*_ = 2.5 and *Τ* = 2 min. Anterior, top; medial, right.

**Movie S8. Actin forms a supracellular cable at the trailing edge of cardioblasts.** Heart development in an embryo expressing *hand-GFP:moesinABD*, a filamentous actin marker. Three stacks covering the length of the heart were acquired every minute and stitched together. Anterior, left; medial, centre.

**Movie S9. The supracellular actin cable resists the backward steps of cardioblasts.** Laser-based tension release in the actin cable at the trailing edge of cardioblasts (top) and contralateral control (bottom) in an embryo expressing *hand:GFP* (green), and *UAS-mCh:moesinABD* driven by *hand-Gal4* (magenta). A stack was acquired every 15 s, and immediately after laser ablation. Time is with respect to the time of ablation. Anterior, left; medial, centre.

